# Plasma exosomal HERV-K transcripts are increased in amyotrophic lateral sclerosis

**DOI:** 10.1101/2025.11.07.686090

**Authors:** Triparna Roy, Misha Ramesh, Nurul Aisha Ahmad Nizam, Steven Tandiono, Khuloud Al-Jamal, Ammar Al-Chalabi, Alfredo Iacoangeli, Ahmad Al Khleifat

**Affiliations:** Department of Biostatistics & Health Informatics, Institute of Psychiatry Psychology & Neuroscience, King’s College London, 16 De Crespigny Park, London, SE5 8AB, UK; Department of Basic and Clinical Neuroscience, Institute of Psychiatry Psychology & Neuroscience, King’s College London, 5 Cutcombe Rd, London, SE5 9RX, UK; Institute of Pharmaceutical Science, King’s College London, London, United Kingdom; Department of Pharmacology and Pharmacy, Li Ka Shing Faculty of Medicine, The University of Hong Kong, Hong Kong, Hong Kong Special Administrative Region, China; King’s College Hospital, London, United Kingdom

**Keywords:** Motor neuron disease, ALS, exosomes, HERV-K, endogenous retrovirus, biomarkers, nanoparticle tracking analysis, neurodegeneration

## Abstract

Human endogenous retrovirus-K (HERV-K) reactivation is increasingly implicated in amyotrophic lateral sclerosis (ALS), with ongoing clinical trials investigating antiretroviral therapies. However, there is limited understanding of how HERV-K is trafficked in peripheral biofluids, and the role of exosomes, nano-sized extracellular *vesicles*, in this process remains largely unexplored. Exosomes offer a stable and cell-specific cargo reservoir that may reflect central pathogenic processes and serve as a minimally invasive biomarker source. In this study, we isolated plasma-derived exosomes from ALS patients (n = 21) and healthy controls (n = 16), and quantified exosomal HERV-K *gag, env*, and *pol* transcript levels using SYBR Green qPCR with RNase treatment and normalization to both traditional and exosome-enriched reference genes. HERV-K *pol* expression was significantly elevated in ALS, with fold-changes ranging from 1.59 to 1.85 (*P* = 0.037–0.051). *env* and *gag* also showed increased expression, though with greater variability. Normalization to the exosome-specific gene *SOD2* provided the most consistent signal. These findings suggest that exosomal HERV-K transcripts, particularly *pol*, could serve as accessible biomarkers for patient stratification and treatment monitoring in HERV-K–targeted ALS trials. This work establishes proof-of-concept for using exosomal cargo to track endogenous retroviral activity in neurodegeneration and supports further investigation of liquid biopsy approaches in ALS precision medicine.

## 1. Introduction

Motor neuron disease (MND), or amyotrophic lateral sclerosis (ALS), is a progressive neurodegenerative disease characterised by degeneration of upper and lower motor neurons, leading to muscle weakness, atrophy, and ultimately respiratory failure (1–4). Despite advances in understanding disease mechanisms, therapeutic options remain limited and confer only modest effects on progression. A major barrier to therapeutic development is the lack of robust, accessible biomarkers that can capture disease-relevant biological processes, enable patient stratification, and provide pharmacodynamic readouts in clinical trials (5–8).

Among emerging molecular pathways implicated in ALS, Human Endogenous Retrovirus K (HERV-K) has attracted increasing attention (8–12). HERV-K belongs to a family of transposable elements integrated into the human genome and retains the capacity for transcriptional activity (9,13–16). Multiple studies have demonstrated increased expression of HERV-K transcripts and proteins in the brain and biofluids of individuals with MND (13,17,18). Clinical studies have linked HERV-K expression to TAR DNA-binding protein 43 (TDP-43) pathology, immune activation, and disease severity, and experimental work has further suggested that the HERV-K envelope (*env*) protein may exert direct neurotoxic effects (9,12,19–21).

This growing body of evidence has led to interest in HERV-K as a potential therapeutic target. Strategies aimed at modulating retroviral activity, including antiretroviral approaches, have been explored in ALS, reflecting a broader hypothesis that aberrant activation of endogenous retroelements may contribute to neurodegeneration (8,22–24). However, translation of these approaches into effective therapies has been limited. One key challenge is the absence of reliable biomarkers that can quantify HERV-K activity in living patients and allow assessment of target engagement and treatment response (18,25).

Extracellular vesicles, particularly exosomes, offer a potential solution to this problem. Exosomes are nanoscale vesicles released into biofluids that carry proteins, lipids, and nucleic acids reflective of their cells of origin (26–28). Their stability in circulation and accessibility from peripheral blood make them attractive candidates for minimally invasive biomarker development (29). In neurodegenerative diseases such as Alzheimer’s disease and Parkinson’s disease, exosome-associated cargo has been shown to capture disease-relevant molecular changes (30,31). In ALS, alterations in exosome size, concentration, and protein content have been reported, including signals related to neurofilament light chain and TDP-43 (28,32–37).

Importantly, exosomes can transport viral and retroviral RNA species, raising the possibility that they may carry HERV-K–derived transcripts (38). This creates an opportunity to detect disease-relevant retroviral activity through a peripheral, cell-derived compartment, rather than relying on bulk plasma or tissue measurements. Such an approach may provide a more specific, quantifiable and biologically informative readout of HERV-K activity, with potential applications in both biomarker development and therapeutic monitoring.

However, despite this rationale, the relationship between exosomes and HERV-K in ALS remains poorly defined. Few studies have systematically examined HERV-K transcripts within exosome-derived RNA. Methodological variability presents an additional challenge (39,40). Exosome isolation techniques differ in yield and purity, RNA content is limited and susceptible to contamination from extracellular sources, and the choice of normalisation strategy can substantially influence gene expression results (28,33,41–43). These factors complicate interpretation and may contribute to inconsistencies across studies.

Given the increasing interest in HERV-K as a therapeutic target, there is a clear need for approaches that can reliably detect and quantify HERV-K activity in accessible patient samples. Exosome-derived RNA represents a promising but underexplored source for such biomarkers, provided that analytical challenges can be addressed.

In this study, we analysed plasma-derived exosomes from patients with ALS and healthy controls to quantify HERV-K *gag, env*, and *pol* transcripts. RNase treatment was used to enrich for vesicle-associated RNA, and multiple reference genes were evaluated to assess the impact of normalisation strategies. We aimed to determine whether exosomal HERV-K transcripts constitute a detectable peripheral signal in ALS and to explore their potential as biomarkers in the context of emerging HERV-K–targeted therapies.

## 2. Materials and Methods

### 2.1. Participant Recruitment and Sample Processing

Plasma samples were obtained from individuals with ALS (n=21) and age and sex matched healthy controls (n=16) through the King’s MND Care and Research Centre, with collection coordinated by the King’s College London MND Biobank.

ALS diagnosis was made according to the revised Gold Coast criteria. Participants were aged ≥18 years and provided written informed consent. Individuals with coexisting neurodegenerative disease, active systemic illness (including infection or cancer), autoimmune disorders, pregnancy, or incomplete clinical data were excluded to minimise confounding factors associated with HERV-K activation.

Blood was collected in EDTA tubes, centrifuged at 2000 × g for 10 min at 4°C, and produced plasma aliquots were stored at −80°C for long term storage.

### 2.2 Exosome isolation, characterisation, and processing

Plasma (300 µL) was centrifuged at 10 000 × g for 15 min at 4°C to remove debris. Supernatants were diluted 1:1 in PBS and ultracentrifuged twice at 130 000 × g for 90 min at 4°C (Beckman Optima MAX-XP, TLA-55 rotor). Pellets were washed in PBS, re-ultracentrifuged under identical conditions, resuspended in 300 µL PBS, and stored at −80°C.

Exosome size distribution and concentration were assessed by nanoparticle tracking analysis (NanoSight LM10; Malvern Panalytical). Samples were diluted in filtered PBS and analysed in triplicates at room temperature using standardised instrument settings.

Dot blot analysis was performed following a previously published protocol (44). Briefly, 40 µl of isolated exosomes (5 x 10^10^ particle/ml) were spotted onto a nitrocellulose membrane (Bio-Rad, UK) and allowed to dry under a gentle stream of nitrogen. The membrane was then blocked with 3% (w/v) milk in Tris-buffered saline containing 0.1% Tween-20 (TBS-T) for 1 hr at room temperature. Following blocking, the membrane was incubated overnight at 4°C with primary antibodies against CD9, CD63 and CD81 (1:1,000 dilution in fresh blocking buffer). The membrane was subsequently washed three times with TBS-T (total wash time of 15 min). The membrane was then incubated with horseradish peroxidase (HRP)-conjugated secondary antibodies (anti-mouse, 1:20,000; anti-rabbit, 1:1,000) for 1 hr at room temperature. After washing as described above, signals were developed using SuperSignal West Femto Maximum Sensitivity Substrate (Thermo Fisher Scientific, UK). Chemiluminescent signals were captured using the Gel Doc imaging system (Bio-Rad, UK) and image analysis was performed using Image Lab software (Bio-Rad, UK).

For protein analysis, vesicles were lysed in 5× RIPA buffer with protease and phosphatase inhibitors on ice for 20 min, followed by centrifugation at 16 000 × g for 5 min. Protein concentration was measured using a bicinchoninic acid assay.

### 2.3 RNA extraction, RNase treatment, and cDNA synthesis

RNA was extracted using the RNeasy Plus Micro Kit (QIAGEN), including gDNA eliminator columns, with additional DNase digestion (20 min at room temperature) to remove residual genomic DNA. RNA yield and purity were assessed by spectrophotometry (NanoDrop). RNase treatment was applied to a subset of samples before RNA extraction to reduce extracellular RNA contamination.

cDNA was synthesised using the QuantiTect Reverse Transcription Kit (QIAGEN) with random hexamers and HERV-K–specific primers, according to the manufacturer’s instructions. cDNA yield and purity were assessed by spectrophotometry (NanoDrop). Amplification specificity and absence of genomic DNA contamination were confirmed by melt curve analysis and 2% agarose gel electrophoresis.

### 2.4 Quantitative PCR (qPCR)

qPCR was performed using PowerUp SYBR Green Master Mix (Thermo Fisher Scientific) and 5 µL cDNA on a QuantStudio 7 (Applied Biosystems). Eight primer sets targeted the HERV-K *pol* region and one set of each HERV-K *gag* and *env* were used to check consistency of gene expression. Housekeeping genes (*GAPDH, ACTB, YWHAZ, SDHA*) and the exosome-specific gene *SOD2* exon 3 were used for normalization. Primer sequences for each gene were created using ERVmap and Primer-Blast (Supplementary Table S1).

Relative gene expression was calculated using the 2^□ΔΔCt^ method, with the mean ΔCt value of the control group used as the calibrator and set to a relative expression value of 1. Relative gene expression was calculated using the 2^−ΔΔCt^ method, with control group samples set to a value of 1 for baseline normalization (45). To ensure accurate normalization, multiple validated housekeeping genes were employed, allowing robust quantification across samples (46,47). ΔCt values were obtained by subtracting the Ct of the housekeeping gene from the Ct of the target gene, and ΔΔCt values were calculated relative to the mean ΔCt of the control group. Agarose gel electrophoresis (2%) was used post-qPCR to visualize amplified products. Melt curve analysis was conducted to confirm specificity.

### 2.5 Statistical Analysis

Statistical analyses were performed using R (version 4.3.1). Group comparisons between ALS (n=21) and controls (n=16) were performed using Mann–Whitney U tests for exosome concentration and gene expression. Comparisons across multiple targets were assessed using Kruskal–Wallis tests. Subgroup analyses by site of onset (spinal n=16; bulbar n=5) were performed using Mann–Whitney U tests. Gene expression analyses were based on values normalised to multiple reference genes.

Multivariate analyses included permutational multivariate analysis of variance (PERMANOVA) and principal component analysis (PCA). Multiple testing was controlled using the Benjamini–Hochberg false discovery rate. Statistical significance was defined as P<0·05.

## 3. Results

Patients with ALS and healthy controls were age and sex matched, and participant demographics are summarised in Table 1. Two participants met criteria for primary lateral sclerosis and were analysed within the ALS group.

**Table 1.** Demographic and clinical characteristics of participants included in the study. The table summarizes the number of ALS patients and healthy controls, including sex distribution, age range, median age, and clinical onset subtypes among ALS cases. PLS cases are listed separately from classical limb- or bulbar-onset ALS.

| Group | N | Female (n) | Male (n) | Age Range (years) | Median Age (years) | Onset Type (n) |
| --- | --- | --- | --- | --- | --- | --- |
| ALS Patients | 21 | 12 | 9 | 35 – 79 | 65 | Spinal Onset: 14<br>Bulbar Onset: 5<br>PLS: 2 |
| Healthy Controls | 16 | 9 | 7 | 30 – 79 | 61.5 | Not Applicable |

### 3.1 Exosome Characterization and Concentration Profiles

Extracellular vesicles from ALS samples showed a broader size distribution with larger median particle size than controls (132 nm [IQR 118–160] vs 104 nm [92–118]; Table 2; Figure 1). Peak particle concentration occurred at ∼100 nm in both groups. Peak concentration was higher in ALS (1·05 × 10^1^□ particles/mL) than controls (0·82 × 10^1^□ particles/mL).

**Table 2.** Median size and concentration of exosomes in ALS patients vs. Controls.

| Group | Median Particle Size (nm) | Size Range (nm) | Median Concentration (particles/mL) | P-value (Median size) | P-value (Total concentration) | P-value (100-300 nm Concentration) |
| --- | --- | --- | --- | --- | --- | --- |
| Controls | 104 | 50–200 | $5.4 \times 10^4$ | 0.015 | 0.02 | 0.018 |
| ALS | 132 | 80–300 | $9.5 \times 10^4$ | | | |

**Figure 1.**
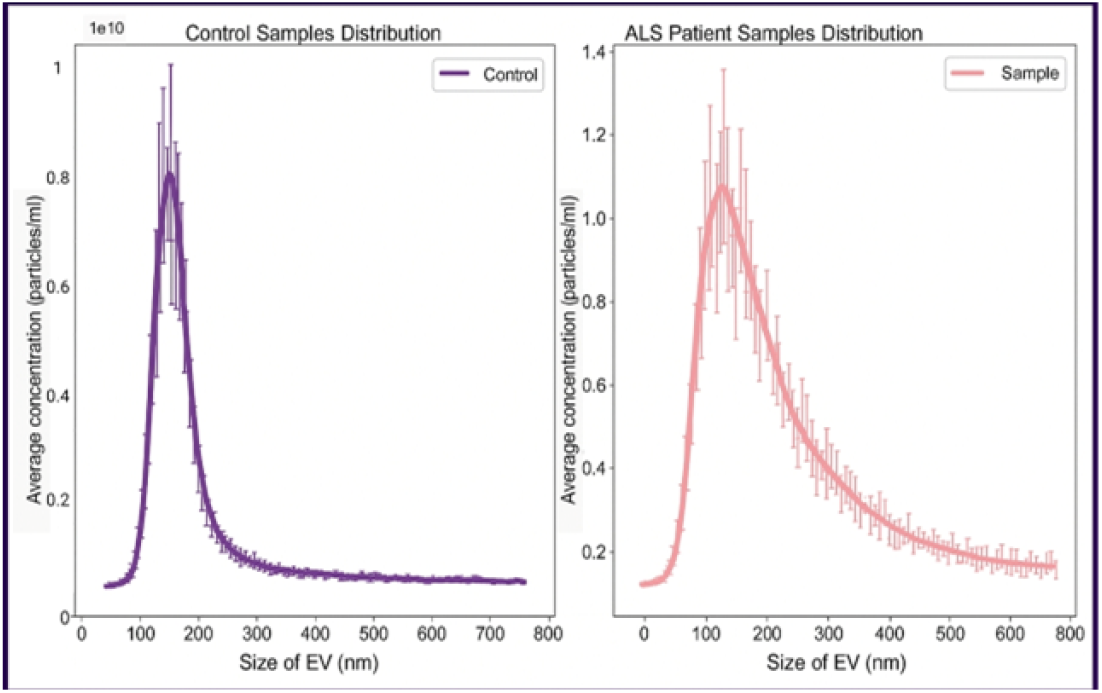
NTA of plasma-derived exosomes in ALS patients and healthy controls. Controls showed a sharp peak at ∼100 nm, while ALS samples exhibited a broader distribution from 80–160 nm and extended toward 300 nm. Median EV size was larger in ALS (132 nm, IQR: 118–160 nm) than controls (104 nm, IQR: 92–118 nm), with significantly higher concentrations within the 100–300 nm size range (*P* = 0.018) and overall concentration differences (*P* = 0.02).

EV concentration in the 100–300 nm range was higher in ALS than controls (median 4·3 × 10□ vs 2·1 × 10□ particles/mL; P=0·018), with a significant difference in overall concentration within this range (P=0·02).

Dot blot analysis confirmed that all exosome samples were positive for the tetraspanin markers CD9, CD63 and CD81 (Supplementary Figure S1), supporting successful isolation and enrichment of exosomes. Variability in signal intensity was observed between samples.

### 3.2 Protein and RNA quality

Bicinchoninic acid assay showed mean protein concentration to be higher in ALS than controls (185·4 µg/mL vs 162·8 µg/mL) (Supplementary Figure S2).

RNase treatment increased RNA yield and purity in both groups. In ALS samples, RNA yield rose from 730.2 ± 13.0 to 805.3 ± 19.2 ng/µL, and in controls from 726.6 ± 12.1 to 800.0 ± 18.7 ng/µL. A260/A280 ratios improved from 1.64 to 1.79 (ALS) and 1.65 to 1.78 (controls), with corresponding increases in A260/A230 (ALS: 1.40 to 1.69; controls: 1.43 to 1.73). DNA yield was also higher following RNase treatment. RNA integrity was confirmed electrophoretically (Supplementary Figure S3), with summary metrics in Supplementary Table S2.

cDNA yields were comparable between ALS and controls but increased with RNase treatment (ALS: 523.5 ± 16.1 to 610.8 ± 21.0 ng/µL; controls: 527.1 ± 15.2 to 613.4 ± 19.9 ng/µL). A260/A280 ratios were correspondingly higher (∼1.78–1.79). Summary cDNA data are shown in Supplementary Table S3.

### 3.3 HERV-K Expression Analysis

Residuals were non-normally distributed across all gene targets (Shapiro–Wilk and Kolmogorov–Smirnov tests, P<0·05), and variance differed between groups for at least one normalisation method (Levene’s P=0·041; Bartlett’s P=0·037). Data transformation (log and square root) was evaluated but did not restore normality or variance homogeneity. Accordingly, non-parametric tests were used for group comparisons.

Relative expression of HERV-K *gag, env*, and *pol* was expressed as fold-change relative to controls (set to 1) and normalised to five reference genes.

Across normalisation strategies, HERV-K transcript expression was higher in ALS-derived exosomes than in controls (Figure 2), although effect size varied by reference gene. Most comparisons reached statistical significance, with weaker separation observed with *SDHA* normalisation.

**Figure 2.**
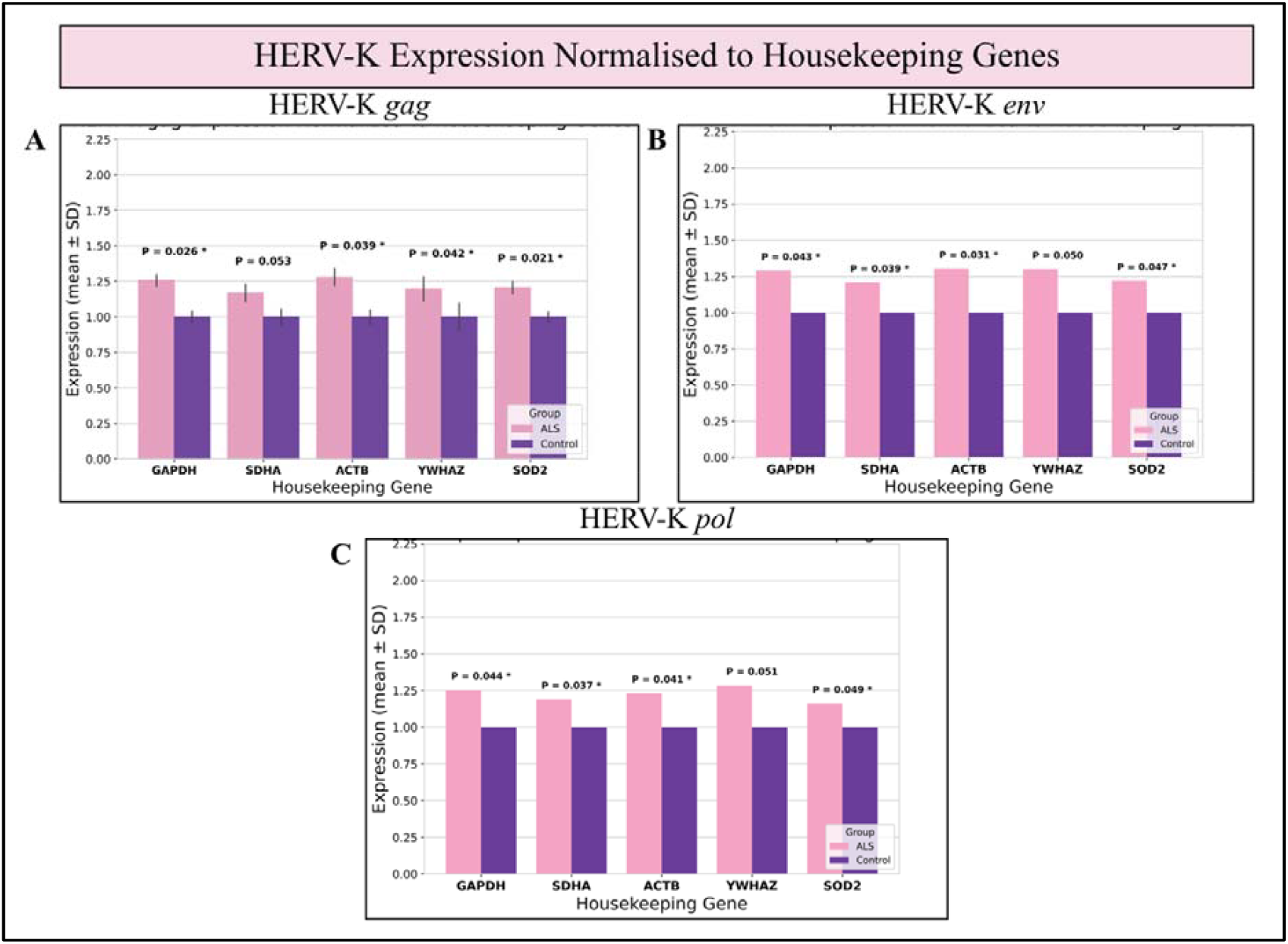
Exosomal HERV-K transcript expression in ALS and controls. Relative expression of HERV-K *gag* (A), *env* (B), and *pol* (C) in plasma-derived exosomes from patients with ALS and healthy controls. Expression was quantified using the 2□ΔΔCt method and normalised to five reference genes (*GAPDH, SDHA, ACTB, YWHAZ, SOD2*), with controls set to 1. Points represent individual samples; bars indicate median values. Group differences were assessed using the Mann–Whitney U test (*P*<0·05). Across normalisation strategies, HERV-K transcripts were consistently increased in ALS, with the most robust upregulation observed for *pol*.

*pol* showed the most consistent upregulation in all three runs across all normalization approaches (fold-change 1.16–1.25; *P* = 0.037–0.049). While *gag* was also elevated, significance was inconsistent across comparisons (*P* = 0.021–0.053). In contrast, *env* expression exhibited higher variability and only marginal statistical support (*P* = 0.031–0.050).

Normalised expression varied by reference gene for HERV-K transcripts (*gag*: H(4)=10·72, P=0·031; *env*: H(4)=10·24, P=0·037; *pol*: H(4)=9·84, P=0·043). Across normalisation strategies, expression was higher in ALS than controls (P=0·021–0·051), with the most consistent differences observed for *pol* (P=0·037–0·049).

Agarose gel electrophoresis confirmed specific amplification, with bands at expected sizes and no evidence of non-specific products (Supplementary Figure S4).

### 3.4 Expression by ALS Site of Onset

HERV-K expression was higher in bulbar-onset (n = 5) compared with spinal-onset ALS (n = 16) across all targets; however, these differences did not reach statistical significance (*gag* P = 0.14; *env* P = 0.095; *pol* P = 0.082; Figure 3). Multivariable regression analyses adjusted for age and sex yielded consistent, but non-significant, positive associations (β = 0.22–0.41; P = 0.091–0.18). Increased variability was observed in the bulbar-onset group, particularly for *pol* expression.

**Figure 3.**
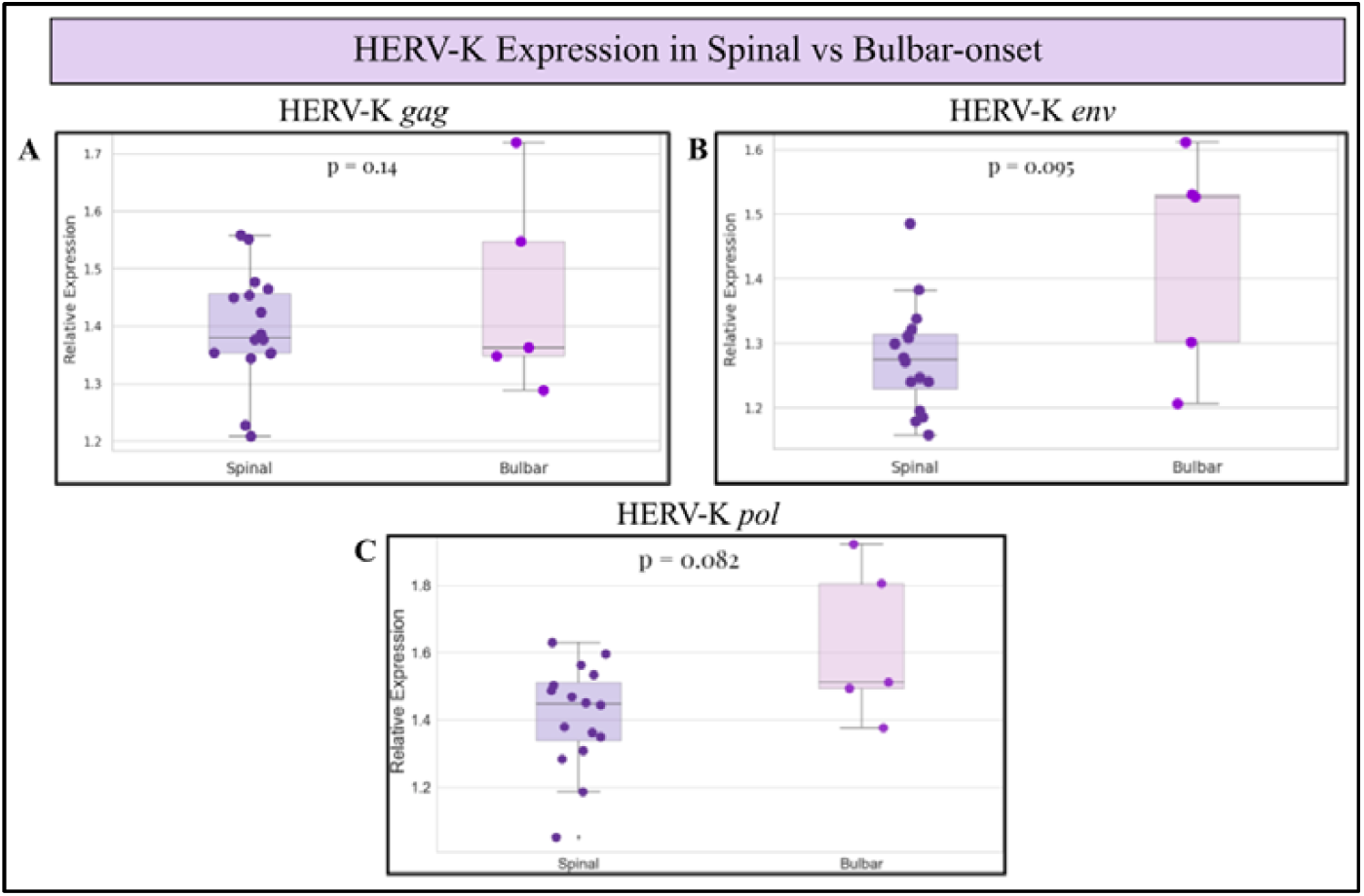
Comparison of HERV-K expression in spinal and bulbar onset ALS. Relative expression of HERV-K *gag* (A), *env* (B), and *pol* (C) comparing spinal-onset and bulbar-onset ALS patients. Across all three viral regions, bulbar-onset patients show a trend toward higher HERV-K expression compared to spinal-onset cases; however, these differences did not reach statistical significance (*gag*: P = 0.14; *env*: P = 0.095; *pol*: P = 0.082; Mann–Whitney U test). Greater variability is observed in the bulbar-onset group.

## 4. Discussion

This study explored whether exosomal HERV-K transcripts differ in ALS and whether such signals might be detectable in peripheral biofluids. Despite comparable EV concentrations, we observed a modest but consistent increase in *gag, env*, and *pol* transcripts in ALS-derived exosomes, with the most reproducible signal observed for *pol*. The concordant increase across all three regions is compatible with a broader upregulation of HERV-K rather than isolated locus-specific effects. This pattern aligns with prior experimental work linking HERV-K expression, particularly *env* to neurotoxicity and immune activation (11,17,24). The relatively stable detection of *pol* across analyses may indicate a more robust marker of retroviral activity, although this requires validation. These findings extend previous evidence of systemic HERV-K activation into the exosomal compartment, suggesting that circulating vesicles may capture aspects of retroviral activity (48,49). However, whether the exosomal signal reflects active pathogenic processes or secondary responses to neuronal injury remains unclear.

Stratified analyses suggested higher *pol* expression in bulbar-onset disease, but this did not reach statistical significance (P=0.082). Although this trend is biologically plausible given the more aggressive clinical phenotype of bulbar-onset ALS, the present study is underpowered to draw firm conclusions. Larger, clinically stratified cohorts will be required to determine whether HERV-K expression contributes to phenotypic heterogeneity.

At the vesicle level, ALS samples were characterised by a shift towards larger extracellular vesicles, rather than clear differences in overall concentration. Whether HERV-K transcripts are actively packaged into these vesicles or represent a downstream consequence of cellular injury remains uncertain. One possibility is that exosomes act as carriers of retroviral RNA, providing a peripheral readout of central disease processes; alternatively, HERV-K expression may represent an epiphenomenon of neurodegeneration. This warrants further investigation into vesicle biology, particularly to determine whether it reflects dysregulated vesicle biogenesis or cellular stress responses.

Another central finding of this study is the sensitivity of results to normalisation strategy. In the exosomal context, where RNA composition differs from total cellular RNA, conventional housekeeping genes may not provide stable reference points. The improved discrimination observed with *SOD2* suggests that compartment-specific controls may better capture vesicle-associated transcripts. At the same time, variability across reference genes highlights an unresolved methodological challenge: the absence of a validated normalisation framework for exosome-derived RNA.

Methodologically, the combination of ultracentrifugation, RNase treatment, and multi-gene normalisation enabled more specific interrogation of vesicle-associated transcripts than approaches based on total plasma RNA. RNase treatment likely improved specificity by reducing extracellular RNA contamination, but introduced variability in downstream cDNA quality, emphasising the need for protocol standardisation. More broadly, reliance on qPCR limits insight into transcript diversity and does not address whether increased RNA translates to functional protein expression.

This study has several limitations. The modest sample size limits statistical power, particularly for subgroup analyses, and increases the risk of both false-positive and false-negative findings. Hence, small effect size, coupled with analytical sensitivity limits the robustness and warrants cautious interpretation. The cross-sectional design further precludes assessment of temporal relationships between exosomal HERV-K expression and disease progression.

Clinical interpretation is also constrained by incomplete metadata, including disease duration, functional status, and treatment exposure, which may have introduced unmeasured confounding. In addition, the absence of disease control groups limits assessment of whether the observed exosomal HERV-K signal is specific to ALS or reflects broader neurodegenerative or inflammatory processes.

At a methodological level, variability in exosome isolation and RNase treatment may have influenced RNA yield and quality, underscoring the need for standardised protocols. The lack of a validated normalisation framework for exosomal RNA remains a key challenge, as results were sensitive to reference gene selection. Furthermore, reliance on qPCR limits resolution of transcript heterogeneity and does not establish whether increased RNA corresponds to protein expression or functional activity.

Finally, although exosome enrichment may improve detection of vesicle-associated transcripts, it remains unclear whether this approach offers meaningful advantages over simpler plasma or serum assays. Establishing analytical robustness, reproducibility, and clinical value will require validation in larger, longitudinal, and well-characterised cohorts.

In conclusion, this study shows that HERV-K transcripts can be detected in plasma-derived exosomes from individuals with amyotrophic lateral sclerosis, with consistent upregulation of *gag, env*, and particularly *pol* compared with controls. These findings extend evidence of HERV-K activation to the exosomal compartment and support the feasibility of using circulating vesicles to capture disease-relevant retroviral signals. Exosomal HERV-K transcripts should therefore be considered an exploratory biomarker (Figure 4). Validation in larger, longitudinal, and clinically stratified cohorts, alongside comparison with non-exosomal approaches, will be required to determine their diagnostic, prognostic, and therapeutic utility in ALS.

**Figure 4.**
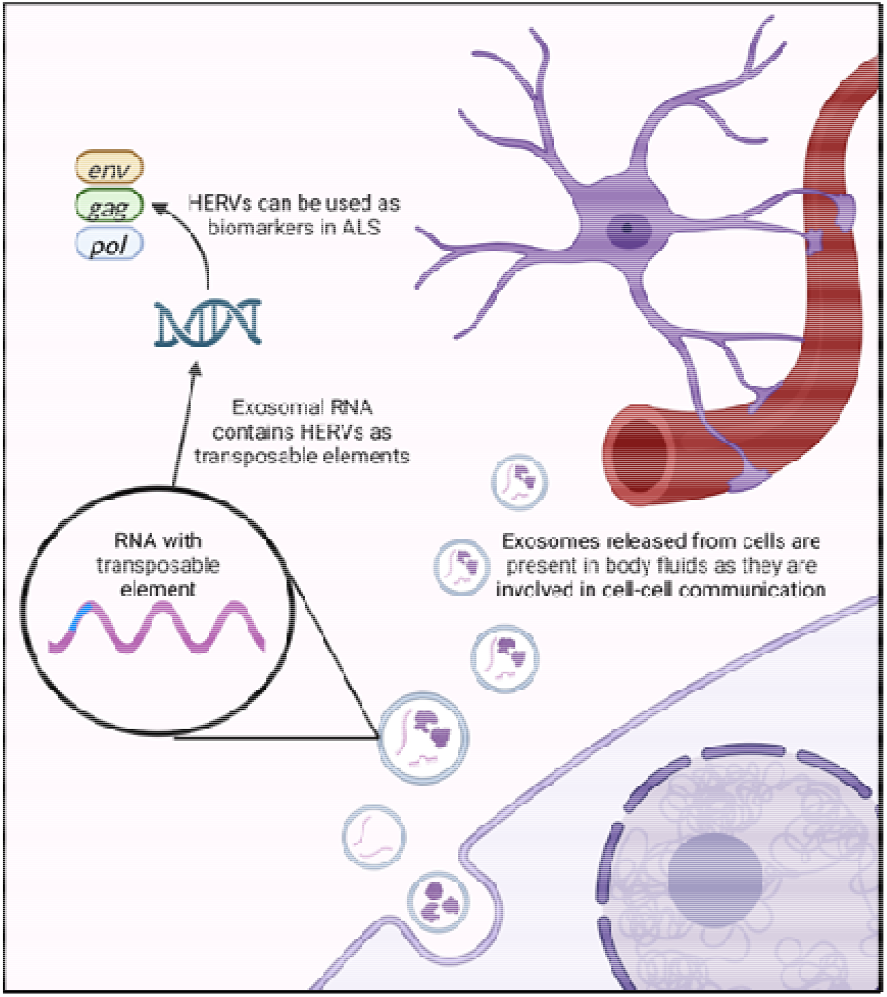
Schematic diagram of exosomes and HERVs as Biomarkers in ALS: Linking Cellular Communication to Neurodegeneration. Created in https://BioRender.com.

## Author Contributions

Conceptualization: TR and AAK; Data curation: TR; Formal analysis: TR and AAK; Funding acquisition: AA-C, AI and AAK; Investigation: TR, AAK and ST; Methodology: TR, MR and AAK; Resources: AA-C and AAK; Supervision: AA-C, AI, AAK and KTA-J; Visualization: TR; Writing – original draft: TR and AAK; Writing – review & editing: AA-C, AI and AAK.

## Funding

TR was supported by the National Institute for Health and Care Research (NIHR) as a Pre-Doctoral Research Fellow (Award Number: 303476) and MND Association. AAK is funded by The Motor Neurone Disease Association (1122462), NIHR Maudsley Biomedical Research Centre, ALS Association Milton Safenowitz Research Fellowship (RE19765), the Darby Rimmer MND Foundation, LifeArc (RE23378), MRC (MR/Z505705/1), and the Dementia Consortium (1819242). AAK is supported by the UK Dementia Research Institute through UK DRI Ltd, principally funded by the Medical Research Council. AI and AAC are funded by South London and Maudsley NHS Foundation Trust, MRC (MR/Z505705/1), MND Scotland, Motor Neurone Disease Association, National Institute for Health and Care Research, Spastic Paraplegia Foundation, Rosetrees Trust, Darby Rimmer MND Foundation, the Medical Research Council (UKRI) and Alzheimer’s Research UK.

## Institutional Review Board Statement

Ethical approval was obtained from the North East – Newcastle & North Tyneside Research Ethics Committee (REC reference: 20/NE/0030).

## Informed Consent Statement

All participants provided informed consent for sample collection and future use in research at the time of enrolment. Newly collected clinical data were anonymised and handled in accordance with GDPR and institutional guidelines.

## Data Availability Statement

Anonymised data may be shared upon reasonable request and under a data-sharing agreement approved by the Chief Investigator. All shared data will comply with the UK General Data Protection Regulation (GDPR, 2018).

## Acknowledgments

The authors acknowledge the KCL MND Biobank for sample provision and thank the MND patient advisory panels. Technical support from the Maurice Wohl Clinical Neuroscience Institute is also gratefully acknowledged. AAC is an NIHR Senior Investigator (NIHR202421) and a Visiting Professor at the Perron Institute for Neurological and Translational Science, Australia. This work was partly supported by an EU Joint Programme - Neurodegenerative Disease Research (JPND) project. The project is supported through the UK MND Research Institute, the following funding organisations under the aegis of JPND - www.jpnd.eu (United Kingdom, Medical Research Council (MR/L501529/1; MR/R024804/1) and Economic and Social Research Council (ES/L008238/1)) and through the Motor Neurone Disease Association, My Name’5 Doddie Foundation, MND Scotland, LifeArc, Alan Davidson Foundation, and Darby Rimmer Foundation. This study represents independent research part funded by the National Institute for Health Research (NIHR) Biomedical Research Centre at South London and Maudsley NHS Foundation Trust and King’s College London. AAK and AI are visiting senior research fellows at the Perron Institute for Neurological and Translational Science, Australia.

## Conflicts of Interest

AA-C declares contracts with the MRC, NIHR and Darby Rimmer Foundation; consulting fees from Amylyx, Apellis, Biogen, Brainstorm, Clene Therapeutics, Cytokinetics, GenieUs, GSK, Lilly, Mitsubishi Tanabe Pharma, Novartis, OrionPharma, Quralis, Sano, and Sanofi). AAK declares contracts with the MRC ((MR/Z505705/1), the Motor Neurone Disease Association (MNDA), National Institute for Health and Care Research (NIHR) Maudsley Biomedical Research Centre, Amyotrophic Lateral Sclerosis (ALS) Association Milton Safenowitz Research Fellowship, Darby Rimmer MND Foundation, LifeArc, and the Dementia Consortium; equipment by NIHR Maudsley Biomedical Research Centre; and consulting fees from the UK National Endowment for Science, Technology and the Arts (NESTA).

## Notes

### Summary of Updates

This revised version incorporates substantial updates to improve methodological transparency, data presentation, and interpretation of findings. The manuscript title has been revised to better reflect the study results. Figure organization has been streamlined, with the conceptual schematic relocated to the end of the Discussion as a graphical summary of the proposed biological framework. Additional extracellular vesicle (EV) characterization data have been included to strengthen validation of EV enrichment. Specifically, dot blot analysis demonstrating the presence of canonical exosome markers (CD9, CD63, and CD81) has been added, together with corresponding methodological details and supplementary figures. Protein quantification (BCA assay), RNA quality assessment, and representative electropherogram data have also been incorporated into the supplementary materials to improve reporting completeness. The Results section has been substantially revised to reduce redundancy and improve clarity. Expression analyses for HERV-K gag, env, and pol transcripts have been consolidated into a summarized presentation, and figures have been reorganized into a unified multi-panel format. Statistical reporting has been corrected and clarified, including refinement of confidence interval interpretation, explicit description of ΔΔCt calculations, and removal of language implying significance where results did not meet statistical thresholds. Additional clarification regarding the inclusion of primary lateral sclerosis (PLS) cases has been added. The Discussion and Limitations sections have been expanded to provide a more balanced interpretation of the findings, including consideration of the clinical utility of exosome-based biomarkers relative to simpler plasma- or serum-based assays. Limitations relating to sample size, lack of disease controls, absence of transmission electron microscopy, and the need for external validation are now discussed in greater detail. Finally, references have been comprehensively reviewed and corrected to ensure that all citations accurately support the associated statements. Collectively, these revisions improve the rigor, transparency, and interpretability of the study while maintaining the central finding that exosome-associated HERV-K transcripts, particularly HERV-K pol, are elevated in ALS and warrant further investigation as candidate biomarkers.

