## Supplementary material for "Plasma exosomal HERV-K transcripts are increased in amyotrophic lateral sclerosis"

### **Supplementary files**

**Abbreviations**

The following abbreviations are used in this manuscript:

| **Abbreviation** | **Full Term** |
| --- | --- |
| ALS | Amyotrophic Lateral Sclerosis |
| MND | Motor Neuron Disease |
| PLS | Primary Lateral Sclerosis |
| PMA | Progressive Muscular Atrophy |
| HERV-K | Human Endogenous Retrovirus K |
| *gag*, *env*, *pol* | Retroviral gene regions encoding core (*gag*), envelope (*env*), and polymerase (*pol*) proteins |
| EV | Extracellular Vesicle |
| NTA | Nanoparticle Tracking Analysis |
| RNase | Ribonuclease |
| cDNA | Complementary DNA |
| qPCR | Quantitative Polymerase Chain Reaction |
| PBS | Phosphate-Buffered Saline |
| RIPA | Radioimmunoprecipitation Assay |
| DNA | Deoxyribonucleic Acid |
| RNA | Ribonucleic Acid |
| *SOD2* | Superoxide Dismutase 2 (exosome-enriched housekeeping gene) |
| *GAPDH* | Glyceraldehyde 3-Phosphate Dehydrogenase |
| *ACTB* | Beta-Actin |
| *SDHA* | Succinate Dehydrogenase Complex Subunit A |
| *YWHAZ* | Tyrosine 3-Monooxygenase/Tryptophan 5-Monooxygenase Activation Protein Zeta |
| *HPRT1* | Hypoxanthine-Guanine Phosphoribosyltransferase 1 |
| ALSFRS-R | ALS Functional Rating Scale-Revised |
| PCA | Principal Component Analysis |
| PERMANOVA | Permutational Multivariate Analysis of Varianc |
| GDPR | General Data Protection Regulation |
| EDTA | Ethylenediaminetetraacetic Acid |
| DNase | Deoxyribonuclease |
| MVB | Multivesicular Body |
| NHS | National Health Service |
| NIHR | National Institute for Health and Care Research |

| **Gene/Gene Region** | **Forward Sequence** | **Reverse Sequence** |
| --- | --- | --- |
| ***GAPDH*** | ACAGTTGCCATGTAGACC | TTTTTGGTTGAGCACAGG |
| ***YWHAZ*** | AACTTGACATTGTGGACATC | AAAACTATTTGTGGGACAGC |
| ***ACTB*** | GACGACATGGAGAAAATCTG | ATGATCTGGGTCATCTTCTC |
| ***SDHA*** | AGCATGCAGAAGTCAATG | ATTTTCCCACAACCTTCTTG |
| ***SOD2* exon3 (Exosome specific)** | CCAGGTGTCGCATTCTGATGTTG | CAGTAGAGCATCTCTCCCAAATG |
| **HERV- K *gag*** | AGCAGGTCAGGTGCCTGTAACAT | TGGTGCCGTAGGATTAAGTCTCCT |
| **HERV- K *env*** | CTGAGGCAATTGCAGGAGTT | GCTGTCTCTTCGGAGCTGTT |
| **HERV- K *pol*** | CTCGCCTTGGAATTCTCCTGTGTTTGT | CAGTGGAAAGTGTTACCTCAGGGAATG |
| **HERV- K *pol*** | GCCTTCACTTTCGCCTTGAATTCT | GCCACCAGGTTTCAGTGGAAAGTG |
| **HERV- K *pol*** | GTGTTTGTAATTCAGAAGAAATCAGGCAAATGGCGTATG | TTGTCAGACTTTTGTAGGTCGAGCTCTTCAACCAGTTAGAGAAAA |
| **HERV- K *pol*** | AACCTATGGGGGCTCTCCAA | TAAGCATTCCCTGGGCAACA |
| **HERV- K *pol*** | ATCAGGCAGATGACGCATGC | CATTCCCTGAGGCAACACTT |
| **HERV- K *pol*** | GTTGCCATCCCCAGTCATGA | AGTCCTGGGGATCCAAAGGA |
| **HERV- K *pol*** | CGCCGTAATTCAACCCATGG | AACTGGTTGAAGAGCTCGAC |
| **HERV- K *pol*** | TGTGGGTAAATCAGTGGCCG | TTCCACTGAAACCTGGTGGC |

**Supplementary Table S1. Primer sequences used for qPCR analysis of exosome-derived cDNA.**Forward and reverse primers are shown for housekeeping genes (GAPDH, ACTB, YWHAZ, SDHA), the exosome-associated marker SOD2 exon 3, and HERV-K *gag, env*, and *pol* regions. Multiple *pol* primers were included to improve detection sensitivity and specificity. Gene expression was quantified using the 2⁻ΔΔCt method, normalised to multiple reference genes, with controls as calibrator (set to 1). Specificity was confirmed by melt curve analysis and agarose gel electrophoresis.

| **Treatment** | **Group** | **RNA Yield (ng/µL)** | **A260/A280** | **A260/A230** |
| --- | --- | --- | --- | --- |
| No RNase | ALS | 730.2 ± | 1.64 | 1.40 |
| No RNase | Control | 726.6 | 1.65 | 1.43 |
| RNase-treated | ALS | 805.3 | 1.79 | 1.69 |
| RNase-treated | Control | 800.0 | 1.78 | 1.73 |

**Supplementary Table S2. RNA yield and purity metrics.** RNA concentration and spectrophotometric purity ratios (A260/A280 and A260/A230) for ALS and control samples with and without RNase treatment.

| **Treatment** | **Group** | **cDNA Yield (ng/µL)** | **Mean A260/A280** |
| --- | --- | --- | --- |
| No RNase | ALS | 523.5 | 1.64 |
| No RNase | Control | 527.1 | 1.65 |
| RNase-treated | ALS | 610.8 | 1.79 |
| RNase-treated | Control | 613.4 | 1.78 |

**Supplementary Table S3: cDNA yield after reverse transcription.**
Summary of cDNA concentrations measured after reverse transcription for ALS and control samples under RNase-treated and untreated conditions.

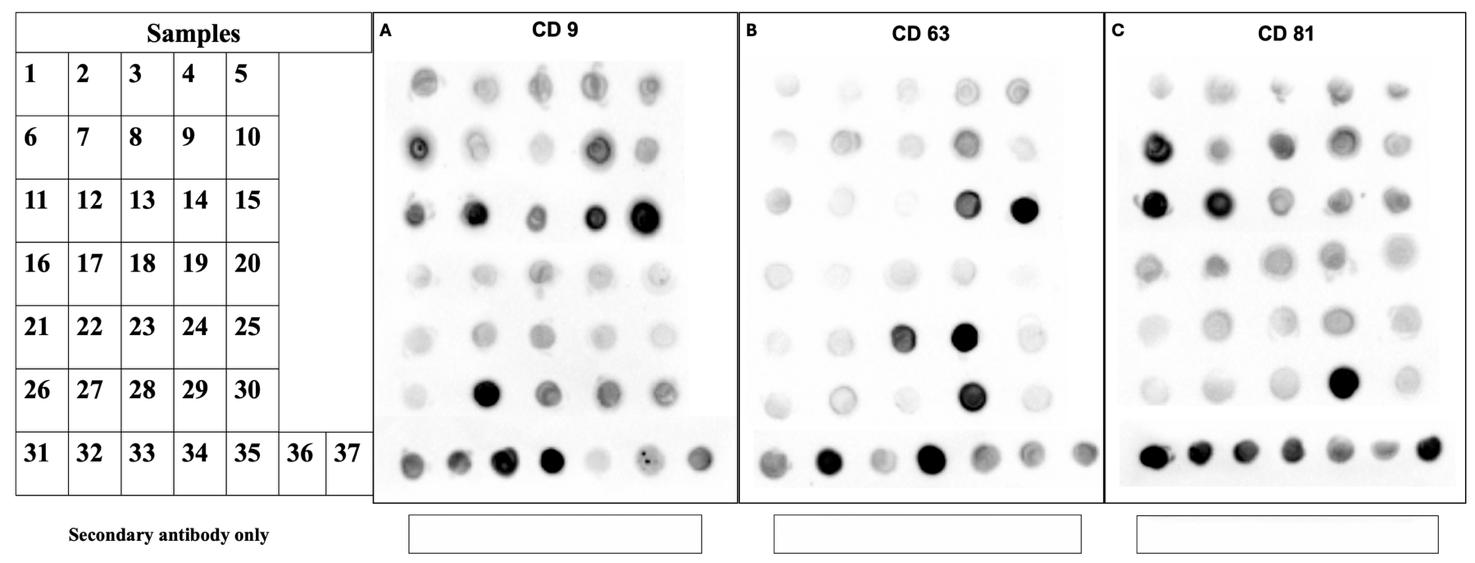

**Supplementary Figure S1. Dot blot validation of exosome surface markers in isolated extracellular vesicles.** Dot blot analysis confirmed the presence of the canonical exosome surface markers **CD9, CD63, and CD81** in ultracentrifuged EV preparations. Each panel shows detection of the respective marker across EV samples (1–37), with a secondary antibody–only control included to assess background signal.

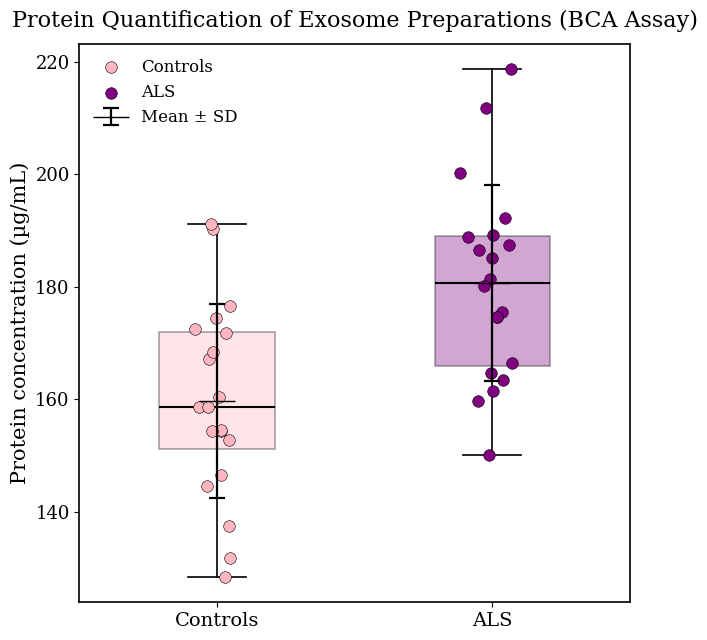

**Supplementary Figure S2: Protein quantification of plasma-derived exosome preparations using the Bicinchonic Acid (BCA) assay.** Protein concentrations of exosome preparations were measured using the BCA assay. ALS samples exhibited slightly higher mean protein concentrations (185.4 µg/mL) than controls (162.8 µg/mL).

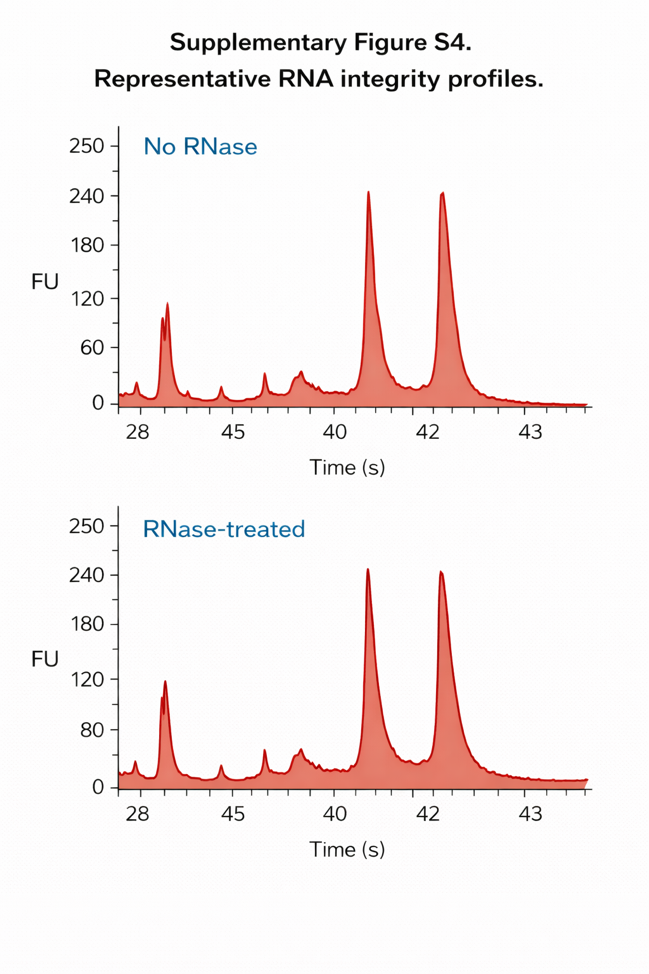

**Supplementary Figure S3. Representative RNA electropherograms.**
Bioanalyzer electropherograms showing RNA fragment profiles of extracellular vesicle RNA samples before and after RNase treatment.

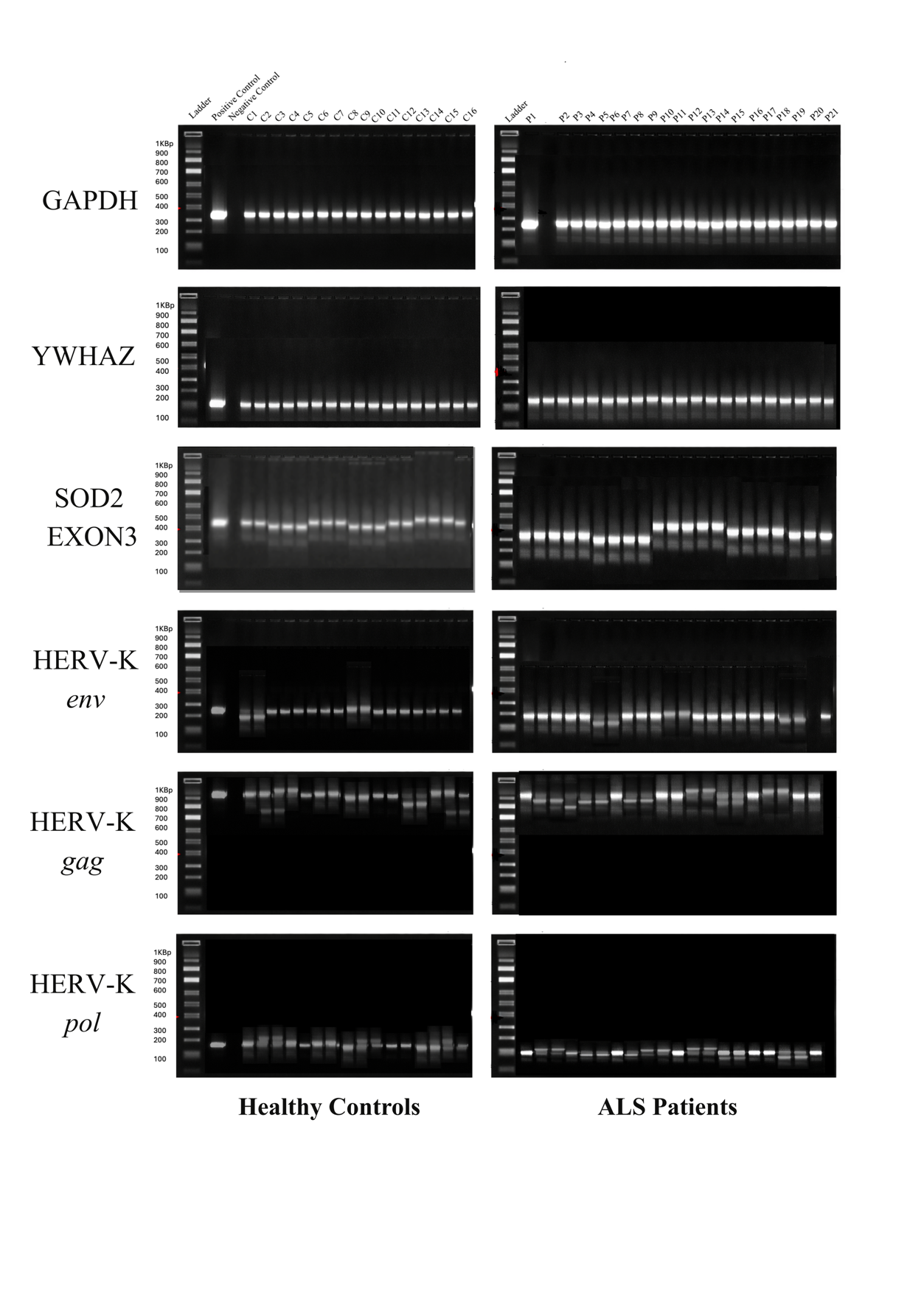

**Supplementary Figure S4. Agarose gel verification of qPCR amplification products for HERV-K genes and reference transcripts.** Representative 2% agarose gel electrophoresis of qPCR amplicons for HERV-K *env* (~256 bp), *gag* (~950 bp), *pol* (~180–190 bp), and reference genes SOD2 (~450 bp), YWHAZ (~250 bp), and GAPDH (~700 bp).
